# Decoding the content of working memory in school-aged children

**DOI:** 10.1101/2023.02.10.527990

**Authors:** Nora Turoman, Prosper Agbesi Fiave, Clélia Zahnd, Megan T. deBettencourt, Evie Vergauwe

## Abstract

Developmental improvements in working memory (WM) maintenance predict many real-world outcomes, including educational attainment. It is thus critical to understand which WM mechanisms support these behavioral improvements, and how WM maintenance strategies might change through development. One challenge is that specific WM neural mechanisms cannot easily be measured behaviorally, especially in a child population. However, new multivariate decoding techniques have been designed, primarily in adult populations, that can sensitively decode the contents of WM. The goal of this study was to deploy multivariate decoding techniques known to decode memory representations in adults to decode the contents of WM in children. We created a simple computerized WM game for children, in which children maintained different categories of information (visual, spatial or verbal). We collected electroencephalography (EEG) data from 20 children (7-12-year-olds) while they played the game. Using Multivariate Pattern Analysis (MVPA) on children’s EEG signals, we reliably decoded the category of the maintained information during the sensory and maintenance period. Across exploratory reliability and validity analyses, we examined the robustness of these results when trained on less data, and how these patterns generalized within individuals throughout the testing session. Furthermore, these results matched theory-based predictions of WM across individuals and across ages. Our proof-of-concept study proposes a direct and age-appropriate potential alternative to exclusively behavioral WM maintenance measures in children. Our study demonstrates the utility of MVPA to measure and track the uninstructed representational content of children’s WM. Future research could use our technique to investigate children’s WM maintenance and strategies.

## Decoding the content of working memory in school-aged children

Working memory (WM), the brain’s limited-capacity system which temporarily maintains information that is no longer physically present (Baddeley & Hitch, 1974; Cowan, 1998; Miller, 1956), has been recognized as the primary determinant of cognitive development in children (for review see Cowan, 2016), and a key predictor of scholastic skills and academic achievement (e.g., Alloway & Alloway, 2010; Bayliss et al., 2003; Bull et al., 2008). It is known that WM performance improves with age (e.g., Gathercole, 1999; Gathercole et al., 2004; Salthouse, 1994), and the emergence of spontaneously used maintenance mechanisms in WM has been proposed as an underlying cause of such improvements (e.g., Camos & Barrouillet, 2011; Gathercole & Adams, 1994; Geier et al., 2009; Magimairaj & Montgomery, 2013; Shimi & Scerif, 2017). However, the way in which some of these maintenance mechanisms are typically measured has made it difficult to build accurate theories of their developmental trajectory. Specifically, children’s WM maintenance mechanisms are usually assessed using behavioral tasks. Yet, the spontaneous use of maintenance mechanisms that occur covertly is difficult to assess behaviorally without introducing specific task manipulations. One concern is that these manipulations can bias whether the to-be-measured maintenance mechanisms can be detected, and how they appear to operate. Such methodological issues can weaken the derivation chain from hypothesis building based on extant theory, through their testing with a given set of methods, to yielding results that are used to build new theory (Meehl, 1990; Scheel et al., 2021). Since science is based on the constant continuation of this cycle, if one element is weak, this puts the inferences that can be drawn from the data at risk on a grand scale. Given the clear educational importance of understanding WM development, methods must be developed which accurately assess how children spontaneously maintain memoranda in WM, i.e., that can uncover and track children’s maintained memoranda without the use of potentially interfering task manipulations. In the current study, we propose an analysis approach using established multivariate analyses of electroencephalographic (EEG) data, known to decode memory representations in adults, to first check whether differences in children’s uninstructed representations can be detected. If successful, this technique could then be used by future studies to investigate questions related to specific maintenance mechanisms and strategies in children.

### Issues in measuring children’s WM maintenance

There are several different strategies that can be used to maintain different types of information in WM for short periods of time (e.g., Bjorklund et al., 2008; Kail & Hagen, 1977). Some of these mechanisms can be measured relatively simply. For example, rehearsal, which involves subvocal repetition of the information to be remembered (Baddeley, 1986; Conrad, 1964) can be tracked by observing lip movements associated with the maintained information (e.g., Elliott et al., 2021; Flavell et al., 1966). Accordingly, there is ample evidence that rehearsal strategies are spontaneously employed without being instructed, from age 7 onwards (e.g., Baddeley et al., 1998; Ferguson et al., 2002; Flavell et al., 1966; Hitch et al., 1991). In another example, children may organize memoranda by a common category during a brief WM delay (Bower, 1970; Mandler, 2002). This organization strategy has typically been measured in children by presenting memoranda in visual forms (e.g., flashcards) that could be spatially grouped by categories (for procedure see e.g., Salatas & Flavell, 1976). Interestingly, only older children (around age 10 onwards) seem to use this strategy spontaneously (Bjorklund & de Marchena, 1984; Hasselhorn, 1992; Schleepen & Jonkman, 2012). However, even 4-year-olds (Sodian et al., 1986), 7-year-olds (Lange & Jackson, 1974), and 9-year-olds (Corsale & Ornstein, 1980) are shown to organize memoranda when task instructions are modified to emphasize the usefulness of the underlying category information. Thus, specific task instructions may alter WM maintenance strategies and influence the measurement of WM maintenance mechanisms. This example foreshadows the complexity of measuring mechanisms that are not directly visible.

Other covert WM mechanisms cannot be easily inferred without employing specific task settings. One such covert mechanism is refreshing, which involves briefly reactivating to-be-remembered information by focusing limited internal attentional resources onto the representation (e.g., Barrouillet et al., 2004; Camos et al., 2018). Though it is not the only covert WM maintenance mechanism in existence, it has received renewed research interest (Káldi & Babarczy, 2021; Lintz & Johnson, 2021; Oberauer & Souza, 2020; Vergauwe et al., 2021; Vergauwe & Langerock, 2023), and is a fitting example mechanism to demonstrate both the necessity to manipulate task parameters to measure it, and how such manipulations can interfere with its measurement. One popular way to measure refreshing is by varying the attentional demands of a secondary processing task (Barrouillet et al., 2009; Bayliss et al., 2003; Bertrand & Camos, 2015; Camos & Barrouillet, 2011; Conlin et al., 2005; Oftinger & Camos, 2015, 2017, 2018; Tam et al., 2010). In such dual-task setups, refreshing would be indexed by declines in WM performance in the maintenance task as a function of the attentional demands of the processing task (e.g., Barrouillet et al., 2009; Bayliss et al., 2003; Camos & Barrouillet, 2011; Tam et al., 2010). A key question has been when refreshing emerges during development, with multiple studies observing an impact of the attentionally demanding processing task (interpreted as the reliance on refreshing) starting at age 7 and becoming progressively greater with age (Barrouillet et al., 2009; Gaillard, 2011; Portrat et al., 2009). However, as Vergauwe and colleagues (2021) have noted, neither the emergence of spontaneous refreshing at age 7, nor the increases in refreshing efficiency from then onto adolescence are unequivocally supported in the literature. First, the detrimental effects of attentional demands on memory performance which are characteristic for refreshing have been observed even at 4–6 years of age (Bertrand & Camos, 2015; Tam et al., 2010). Second, memory performance decreases as a result of attentionally demanding concurrent tasks have not been found to differ much between 6-year-olds and 8-year-olds (e.g., Conlin et al., 2005; Oftinger & Camos, 2015, 2017, 2018). Though dual-task paradigms have yielded mixed results on the developmental trajectory of refreshing, these paradigms have still all detected refreshing in children. Removing the secondary task, however, seems to make refreshing undetectable. Namely, in a paradigm without a dual task design, and using a different outcome measure than the above studies (based on Vergauwe & Langerock, 2017; see also Vergauwe & Langerock, 2023), Vergauwe and colleagues (2021) did not find evidence for spontaneous refreshing in children older than 7 years of age (see also Valentini & Vergauwe, 2023). However, the removal of the processing task could have had other consequences, for example rendering the paradigm less challenging for children and easier to understand. The results presented here demonstrate that specific paradigm design choices, for example the outcome measure and the inclusion of a secondary task, can influence whether or not refreshing can be detected in children. The contrast between paradigms with and without secondary tasks has the potential to upend common conceptions of WM development and refreshing as a maintenance mechanism. Combined with the results regarding the organization strategy, on a larger scale, these outcomes (as well as e.g., Brady et al., 2021) highlight the need for deeper considerations of methodology before testing theoretical assumptions in the WM field.

### Moving forward: Designing better methods for assessing WM maintenance in children

Most studies on children’s WM mechanisms have used behavioral measures exclusively, meaning that detecting maintenance mechanisms was reliant either on the observable production of said mechanisms (for overt mechanisms), or inferences based on results under specific task settings (for covert mechanisms). Though such approaches have been instrumental to understanding WM development, we have seen that they are limited, in that it may be challenging to disentangle whether failure to detect an effect in children is due to a) a true absence of an effect, b) the children not being motivated to do a difficult task, or c) the children not understanding the instructions of the task. Therefore, it would be powerful to develop a measure that is independent from task demands, which would allow children to behave, and process information spontaneously.

A potential solution lies in departing from behavioral-only measures. Assessing maintenance mechanisms using neural measures would remove the need for introducing secondary tasks or elaborate instructions, thus simplifying and removing sources of bias from behavioral tasks, and allow children to maintain memoranda in whichever way was most natural to them. In particular, Multivariate Pattern Analysis (MVPA) can index different perceptual or cognitive states, by training classifiers to distinguish between different patterns of activity that are distributed across the brain (Haxby et al., 2001; Norman et al., 2006). Multivariate analyses of electroencephalography (EEG) data, has successfully decoded the contents of attended information in WM (LaRocque et al., 2013), the amount of WM load (Adam et al., 2020), individual differences in WM load and attentional focus (Karch et al., 2015), and attentional processes involved in the transfer of information into long-term memory (deBettencourt et al., 2021), all in adults. It has less frequently been applied to neural data collected from children, though there have been some notable exceptions (e.g., Mares et al., 2020 [face processing in typical development]; Petit et al., 2020 [language processing in typical development and autism]; Lui et al., 2021 [word reading skills with Chinese characters]). To our knowledge, however, this method has yet to be used to elucidate WM processes in children, let alone WM maintenance processes in children. With that, it would be prudent for new decoding approaches in children’s WM maintenance to first make sure that basic differences in maintained content are detectable (e.g., as in LaRocque et al., 2013 in adults) before attempting to differentiate between different maintenance mechanisms. To that end, we conducted a proof-of-concept study to check if such basic differences in uninstructed WM representational content are decodable, as a first step towards one day being able to use MVPA to distinguish between different uninstructed maintenance mechanisms in children.

### The current study

In the current study, we aimed to directly assess the uninstructed content of children’s WM during *maintenance* and track its changes over time, using the MVPA technique which is known to decode memory representations in adults. To do so, we posited that, instead of changing behavioral task parameters (i.e., without relying on difficult concurrent processing tasks, or hard-to-understand task instructions), we could leverage measures of brain activity during maintenance. We tested this hypothesis by combining a simple, child-friendly, computerized working memory task with multivariate analyses of electroencephalographic (EEG) measures. Our study design allowed us to probe the uninstructed representational content of children’s working memory during maintenance, and throughout the course of an entire trial from encoding to response. We decoded between three categories that reflect the most commonly used stimulus categories in WM research (visual features, spatial locations, verbal stimuli), mapping onto domain-specific dissociations of working memory resources as proposed in the popular multi-component model (Baddeley & Hitch, 197; Baddeley, 1986; Baddeley & Logie, 1999). The ability to decode the contents of WM from children was the key aim of our study, but we also went on to assess the validity and reliability of our approach through a series of exploratory analyses.

## Methods

### Participants

A total of 25 children (8 female, mean age = 9 years 4 months, SD age = 1 year 3 months, Range: 6 years 0 months – 12 years 9 months) were recruited for the present study, through personal contacts, word of mouth, and the participant database of the Working Memory, Cognition and Development lab. The sample size was determined based on prior studies using MVPA on children’s EEG data available to us at the time (Lui et al., 2021; Mares et al., 2020; Petit et al., 2020). Of this, five participants were excluded due to not finishing the task, having behavioral accuracy at or below chance level (i.e., 50% of the total accuracy), or excessive noise in their EEG data (that could not be cleaned from the data such at least 200 trials remain in the dataset, e.g., due to excessive movement throughout the experimental task). These exclusion criteria were determined before data collection. Thus, the final sample included 20 children (7 female, mean age = 9 years and 7 months, SD age = 1 year and 5months, Range: 7 years 0 months – 12 years and 2 months).

Participants were tested at the EEG lab of the Faculty of Psychology and Educational Sciences of the University of Geneva and were offered a 20 Swiss franc voucher from a popular media store. All research procedures were approved by the University of Geneva Ethical Commission (approval code: CUREG_2021-05-49). Informed consent was obtained from parents/caregivers and verbal assent was obtained from children before participating in the study.

### Stimuli

Stimuli belonged to one of three categories: Visual (an image of a robot), Spatial (an image of a rocket ship in a given spatial location on a circular grid), or Verbal (an image of a French-sounding non-word, i.e. the written form of a French-sounding non-word). There were 16 stimuli per category. For the spatial category, there were 16 positions that a rocket could occupy on a circular display; based on the memory items from Ricker and Vergauwe (2020). For the visual category, there were 16 images of robots (generated by typing the following numbers into the ‘generate’ query space at Robohash.org: 10, 20, 30, 40, 50, 60, 70, 80, 90, 100, 110, 120, 130, 140, 150, 160, and saved as a .png file). Finally, for the verbal category, there were 16 nonwords presented in written form on the screen. The nonwords were written in white letters in Calibri font, size 115. The nonwords were generated by WinWordGen 1.0 (Duyck et al., 2004), had 5 letters and 2 syllables each, and were drawn from Lexique.org with a base language of French (all other settings were set to default). A list of 119 words was generated, and we selected 16 that were verified not to resemble a real French word by a native French speaker (i.e., *karir, gagon, wacha, eurit, hanre, impet, asers, podou, coune, sarda, tindi, lurre, madru, ouail, pelme*, and *usase*). The output of the program and the verified list if words are both included in the OSF repository for this project (see Open Practices Statement below). All stimuli were presented centrally on a black background subtending 5 degrees of visual angle, to minimize eye movements. For comparison purposes, we include an image of all the stimuli together In the Supplementary Materials (Supplemental Figure 1).

### Task procedure

The experimental task involved a single-item delayed-recognition task (Figure 1), based on the Phase 1 task of Experiment 1 in LaRocque et al. (2013). Importantly, participants were only told to remember the information – there were no instructions as to how to maintain the stimuli. Each trial began with an inter-trial interval (ITI, average duration 1000ms), during which participants were presented with a white fixation dot centrally on the screen. To reduce anticipation effects, the ITIs were randomly jittered between 800ms to 1200ms in steps of 50ms. Next, during the Sensory period, the to-be-memorized item was displayed for 1000ms. Stimulus category was randomly determined on each trial. Afterwards, during the maintenance (Delay) period (2000ms), a central white fixation cross appeared on the screen. Then, a probe stimulus of the same category was displayed for 1000ms. Finally, during the Response period, a white question mark was displayed centrally (until the participant responded or 2000ms, whichever was shorter), indicating to the participants to respond. If the probe stimulus matched the to-be-memorized stimulus, participants were to press the ‘k’ key on the keyboard in front of them. If the probe stimulus did not match the previously encoded target stimulus, participants were supposed to press the ‘d’ key on the keyboard. The probe stimulus matched the to-be-memorized stimulus 50% of the time. The ‘k’ and ‘d’ keys were marked with green circular stickers. Prior to the experiment, participants were told to respond when they saw the question mark on the screen. There were no instructions on which hand to use to provide responses. To minimize blinking artefacts during the Sensory (encoding) and Delay (retention) periods, participants were encouraged to withhold blinking during these times, and we turned on the light in the testing booth for participants that had trouble withholding blinking.

**Figure 1.**
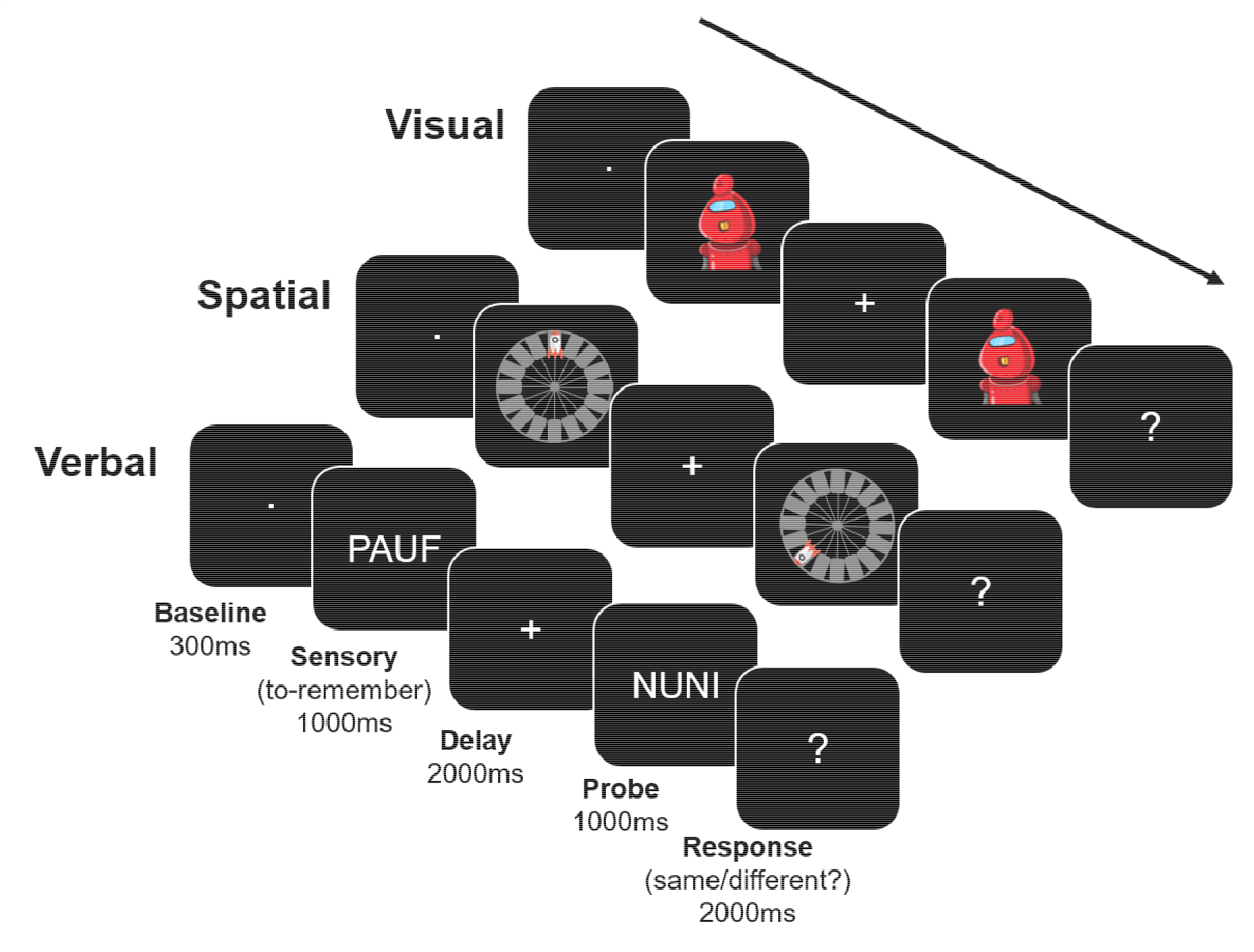
Task schematic. One trial per category (visual, spatial, verbal) is depicted. The stimuli for the visual category were images of robots, the stimuli for the spatial category were rockets in a particular location on a platform, and the stimuli for the verbal category were nonwords. The Baseline period was the last 300ms of the ITI, which lasted 1000ms on average. The stimuli were presented during the Sensory period (1000 ms), followed by a blank Delay period (2000 ms). Then, a probe image appeared (1000 ms) that was either the same image (50% of the time) or a different image from the same category (50% of the time). Finally, during the Response period (2000 ms) a question mark appeared, and participants could respond. Categories were randomly intermixed across trials over the duration of the experiment. Note that the stimuli are shown much larger in the figure, for clarity, than they appeared in the experiment. EEG data were epoched so as to contain the Baseline, Sensory, and Delay periods (i.e., - 300ms to 3000ms relative to Sensory period onset).

Participants completed 128 trials per category, resulting in a total of 384 experimental trials across categories. To help increase children’s motivation, the task contained a background story (helping astronauts find their way home from an alien space base) and was presented in a game-like fashion. Participants could take self-timed breaks after every 48 trials. During these breaks, the participants’ total number of correct responses out of possible correct responses was shown on the screen, alongside the number of ‘blocks’ left. Before starting the paradigm, participants completed several slower practice trials. In total, each session took a maximum of 2h, with approximately 45 minutes of data collection.

The experimental paradigm was programmed using the Psychopy Builder Standalone version 2020.2.5 (Peirce et al., 2019), and presented on a 24” LCD monitor (60Hz refresh rate) in a sound-attenuated, shielded booth. A BioSemi ActiveTwo amplifier (BioSemi Inc., Amsterdam, The Netherlands) was used to record EEG data from a 64-electrode BioSemi gel headcap (10/20 electrode layout). All sites were referenced online to electrode Cz, and re-referenced offline to the average reference. To record eye movements and blinks, additional electrodes were placed at the outer canthi of both eyes (for the horizontal electrooculogram; HEOG) and above and below the right eye (for the vertical electrooculogram; VEOG). Electrode impedances were adjusted to below 5 kΩ prior to the start of the experiment. Data were digitized at 2048 Hz.

### Behavioral data analyses

Although behavioral data analyses were not the focus of the present study, we calculated accuracy (percentage of correct responses) over the entire task. We excluded the data of participants with accuracy that was at or lower than a level that would be obtained by pure chance (50%) from further EEG analyses. To estimate whether our participants could successfully complete the task, we derived the average accuracy score across all participants at three different points: before any exclusions, after exclusions based on behavioral criteria, and after exclusions based on EEG-related criteria (i.e., the data retaining at least 200 trials after cleaning).

### EEG preprocessing

For EEG data preprocessing, we used the Matlab-based (Natick, Massachusetts: The MathWorks Inc) EEGLAB software (v.2022.0, Delorme & Makeig, 2004). We first down-sampled the data to 500Hz, removed the DC offset, and applied a bandpass filter of 1Hz – 40Hz (12 dB/octave roll-off computed forward and backward to eliminate phase shift). Then, we epoched the data from -300ms to 3000ms relative to the onset of the stimulus for each trial, such that each epoch contained the Baseline period (-300ms to 0ms, during the ITI), the Sensory period (0ms – 1000ms), and the Delay period (1000ms – 3000ms; see Figure 1). A semi-automatic artefact rejection procedure was used to remove artefacts (transient noise, movement, skin potentials, etc.), which consisted of applying an automatic artefact rejection criterion of ±150μV for EEG artefacts (adapted to children’s EEG, see e.g., Melinder et al., 2010; Shimi et al., 2015) along with visual inspection. Next, to remove the influence of blinks, we conducted independent component analysis (ICA) using the ICLabel package (Pion-Tonachini et al., 2019) in EEGLAB. We detected those components that contained eye movements and blinks with visual inspection and removed only these components from the data. We discarded any electrodes contaminated by artefacts, based on visual inspection (maximum 13% of the electrode montage) and interpolated the missing data using 3-dimentional splines (Perrin et al., 1987). Our EEG analyses only included participants with over 200 trials (67% of the total number of trials) remaining after the cleaning procedure.

### EEG classification

The main goal of this study was to examine the EEG multivariate representations for children performing a working memory task. We approached this goal in two ways: 1) We calculated the average classification accuracy during the Baseline, Sensory, and Delay periods separately (time-average classification), and 2) We examined how classifier performance trained at a specific moment in time generalized over other timepoints (temporal generalization classification). This set of analyses allowed us to probe whether we can detect differences in the uninstructed representational content of children’s working memory. First, we tested whether we could reliably decode WM content during delay periods, as well as other periods during the trial. Second, we further tracked the representational content over time by showing how long the same representational structure can be detected.

Multivariate classification was performed within each subject by employing the MVPA-Light toolbox (Treder, 2020) with the linear discriminant analysis (LDA) classifier. We used each participant’s preprocessed single-trial voltage amplitudes across all EEG channels as input features for the classifier. Both correct and incorrect response trials were included in the analysis (as in LaRocque et al., 2013), in order to maximize the number of trials per category, assuming that incorrect responses were made because the details of a given stimulus, and not its category membership, were wrongly remembered.

For the time-average classification, we calculated the average voltage amplitudes across each of the respective time-windows: -300ms – 0ms for the Baseline period, 0ms – 1000ms for the Sensory period, and 1000ms – 3000ms for the Delay period. We averaged across trials, with 5 samples for each average and demeaned the data across trials. We performed 100 iterations of the classification analyses for each time-bin (as in Adam et al., 2020). Cross-validation was applied, such that, on each iteration, 2/3 of the trials were randomly assigned to a training set and 1/3 of the trials to a held-out test set. The classifier performance was determined by averaging the classification accuracy across the 100 iterations. To ensure the same number of trials per category in the training and test set, condition labels with fewer trials were up-sampled using a stratification procedure, natively implemented in the MVPA-light toolbox. The resulting mean trial numbers per condition were as follows: 99.95 (SD=16.29) for the visual condition, 101.75 (SD=17.01) for the spatial condition, and 101.65 (SD=15.55) for the verbal condition. Thus, the conditions had very similar trial numbers, making for a rather balanced dataset. The outputs of the classification performance were accuracy (i.e. the proportion of correctly predicted class labels) and a set of confusion matrices (i.e., tables of proportions correctly predicted and incorrectly predicted class labels). Significance was statistically assessed against theoretical chance (33%) via Bonferroni-corrected one-sided t-tests at each time-bin (assuming no meaningful values that are below chance). Statistical assessment against theoretical chance involved performing subject-wise permutation (see also Fahrenfort et al., 2018), with 1000 iterations per subject, at an alpha level of 0.05, implemented via the “*mv_statistics*” function of the MVPA-light toolbox.

To analyze temporal generalizability, we divided each trial into smaller 50-ms time-bins, averaged across 5 trials, and calculated the mean voltage amplitudes for each bin (as in Adam et al., 2020). Then, for each time point, we trained the classifier on recorded brain activity for the given time point and tested it on brain activity recorded at all other time points. Significance was statistically assessed against theoretical chance (33%) via Bonferroni-corrected one-sided t-tests at each time-bin. The results of the 50-ms temporal generalization are reported below. The same analysis was also run using 20-ms time-bins, revealing very similar results (see Supplementary Materials).

To further validate our approach and investigate its robustness, we conducted additional exploratory analyses to assess its reliability and validity. First, we split our EEG dataset in half and conducted several analyses to establish the robustness of EEG decoding within and across these halves. Specifically, for each participant, we split the data into an early half (blocks 1 – 4 in the experimental session) and a late half (blocks 5 – 8), and applied the same temporal generalization procedures to each half separately. Afterwards, we cross-decoded from the early to the late half, such that data from the early half of the session were used for training the classifier and the data from the late half of the session were used for testing the classifier (see e.g., LaRocque et al., 2013; Lewis-Peacock et al., 2012). Second, we descriptively compared the results of the main time-average decoding to an expected pattern based on our given task design (i.e., no above-chance decoding at Baseline, high above-chance decoding at Sensory, and lower but still above-chance decoding at Delay). Third, we descriptively compared the results of the confusion matrix generated by the time-average classification above to a pattern that follows classic WM theory (i.e., that verbal stimuli would be the least confusable with other categories, while visual and spatial stimuli would be more confusable with each other, though still distinct). Fourth, we investigated the relationship between decoding accuracy in the Sensory and Delay periods per individual, to capture the consistency of our measures across participants. Here, we had two assumptions: 1) that classification performance when observing the stimulus to be remembered (i.e., during the Sensory period) should be higher than classification performance when maintaining said stimulus in memory in its absence (i.e., during the Delay), and 2) that individuals with higher classification performance at Sensory than others will also have higher classification performance at Delay than others. To test these assumptions, we first compared average classification accuracy at Sensory and Delay using a paired-sample right-tailed t-test. Then, we quantified the strength of the relationship between classification accuracy at Sensory and Delay via Pearson correlation. Finally, we examined decoding accuracy as a function of age. We did so first by using a linear regression between participants’ age and their average classification accuracy for the Baseline, Sensory, and Delay periods separately, using the “fitlm” and “anova” functions in Matlab. This was done to rule out that our results were driven by older participants, and not the entire range of participants. It is well-established that children’s WM performance tends to improve with age (e.g., Gathercole, 1999; Gathercole et al., 2004; Salthouse, 1994), and thus our above-chance decoding results could have stemmed from older ‘high performers’ with higher decoding accuracy. Finally, we compared behavioral accuracy and classification accuracy between the youngest participants in the sample and the entire sample. Since our youngest participants were between 7 and 8 years of age (5 participants aged: 7 years, 7 years and 6 months, 7 years and 9 months, 8 years and 1 months, and 8 years and 3 months), they might have had trouble processing the nonwords due to less efficient reading skills. To ascertain if decoding results were driven by such a difference in processing verbal and other stimuli in young participants, we compared their behavioral and classification performance across categories to that of the entire sample. Behavioral data were analyzed using a Bayesian analysis of covariance (BANCOVA) with one within-subjects factor of Category and Age as a covariate, using the default settings (Cauchy prior with a 0.707 scaling constant for effect size) in JASP v.0.16.4.0 (JASP Team, 2020). We also descriptively compared the youngest participants’ accuracies per category with those of the entire sample. For the decoding analyses, we extracted a confusion matrix for the youngest participants only, using the same methods as for the main analyses described above, and compared the values per category with those of the entire sample. To avoid reiterating the main analysis results, in the Results section, we will only present the results of the split-half analyses, confusion matrix analysis, individual difference analyses, and analysis of decoding accuracy by age.

### Open Practices Statement

The behavioral data, exclusion criteria, preprocessing, analysis scripts, and study materials (stimuli and task design) are available at https://osf.io/jeh67/?view_only=2a9c2379a1514ce996b446cf1b0690b3. Due to space limitations on OSF, the raw EEG data are stored in a Zenodo repository: (https://zenodo.org/record/6555702 [until publication, email fist author for access]), and the preprocessed EEG data are stored in another Zenodo repository: (https://zenodo.org/record/6580005 [until publication, email fist author for access]). No part of the study procedures or analyses was pre-registered prior to the research being conducted. We report how we determined our sample size, all data exclusions, all inclusion/exclusion criteria, whether inclusion/exclusion criteria were established prior to data analysis, all manipulations, and all measures in the study in the pages above.

## Results

### Behavioral results

In a first analysis step, we verified whether participants successfully performed the task by checking their average behavioral accuracy results. The total sample had an average accuracy score of 84% (SD = 21%). After applying our behavioral exclusion criteria, the average accuracy score was 89% (SD = 9%). Finally, after applying our EEG exclusion criteria, the average accuracy score was 90% (SD = 8%). This high accuracy demonstrates that our participants successfully performed the WM task.

### EEG classification

The key goal of this study was to use multivariate pattern analysis to decode EEG data collected from children during a WM game. We were interested in whether multivariate EEG patterns differed when children maintained different categories of information (visual, spatial, and verbal). First, we examined the time-average EEG voltage patterns during the Baseline, Sensory, and Delay periods (Figure 2). As expected, we found that decoding accuracy was above theoretical chance (33%) in the Sensory period (mean = 58.0%, median = 60.0%, *SD* = 11.1%, p = 2.77e^-09^) and the Delay period (mean = 44.4%, median = 45.1%, *SD* = 8.6%, p = 7.38e^-06^), but not in the Baseline period (mean = 34.6%, median = 33.8%, *SD* = 4.4%, p = 0.11). This demonstrates reliable decoding of children’s WM contents during the time when children maintained information (Delay), and when they viewed information (Sensory), but also verifies that we were unable to decode during the moments prior to the stimulus presentation when they were preparing for the onset of the next trial (Baseline).

**Figure 2.**
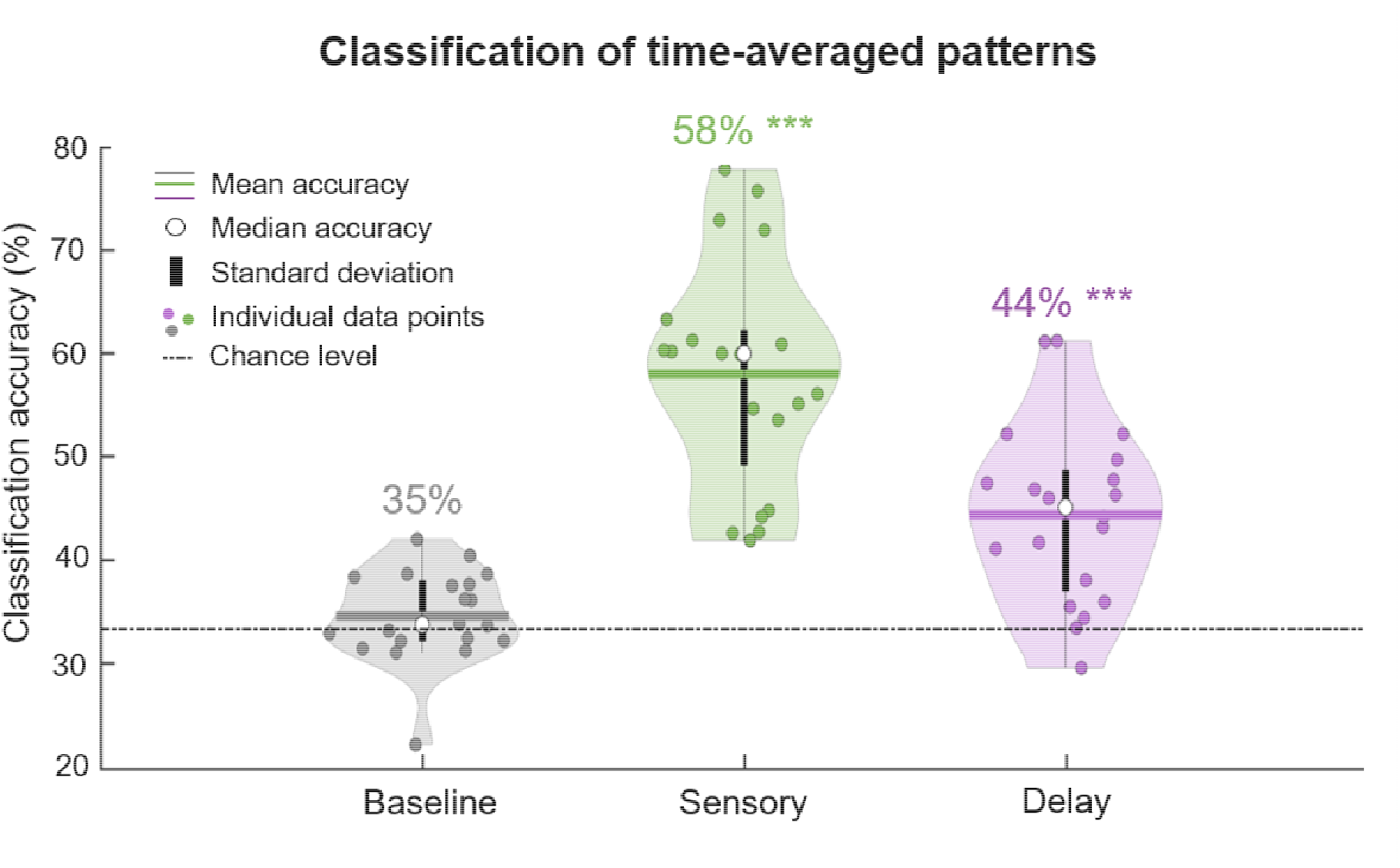
Classification accuracy of time-averaged patterns during the Baseline (gray), Sensory (green), and Delay (purple) periods. The mean is illustrated as a horizontal line, the median accuracy is depicted as a white circle, and standard error is depicted as a vertical black line. Individual data points are depicted as dots, and theoretical chance is represented by a dashed and dotted horizontal black line. The numerical average accuracy score is displayed above each violin plot. Accuracy scores that are significantly above chance are marked with *** if they are above the *p*<0.001 threshold.

### Temporal generalization of EEG patterns

Next, we examined how children’s WM representations unfolded across time by examining how patterns generalized across different moments (Figure 3). That is, we trained a multivariate classifier at a particular time point, and tested it on all other timepoints. We observed similar multivariate representations across time, particularly from the Sensory to the Delay periods of the trials: Representations that were detected during the early Sensory period could be traced with high accuracy over the entire Sensory period, and importantly, for a portion of the Delay period. This is reflected in reliable decoding at distant moments of time, far from the diagonal of the matrix. In sum, our results suggest not only that differences in observed and maintained representational content can be detected, but that the same representations that were formed when stimuli were observed persisted in WM for a portion of the maintenance period, even when the stimuli were absent.

**Figure 3.**
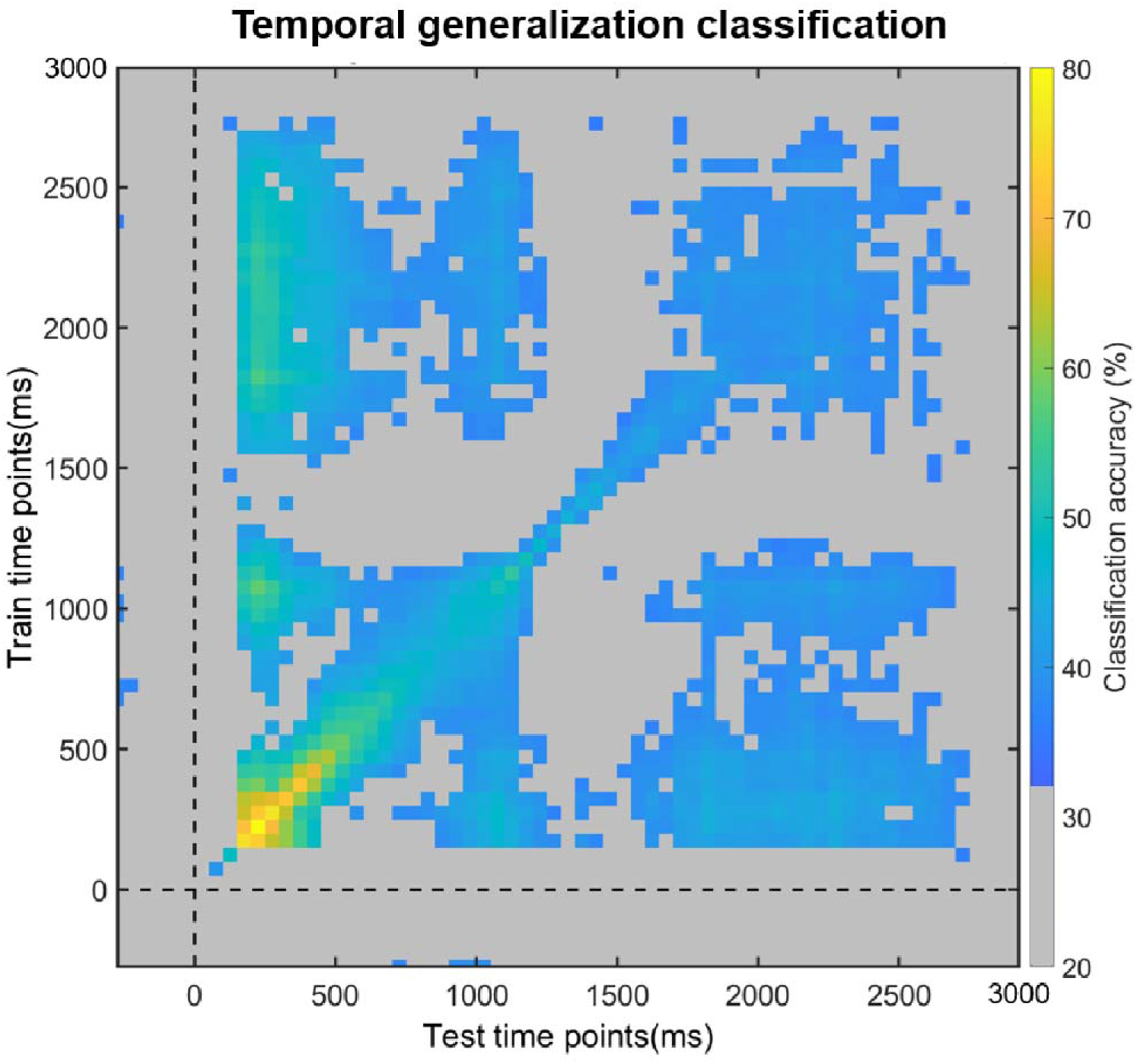
Temporal generalization of classification, where a classifier was trained on one time point and tested on all other time points. Average classification values are overlaid by a significance mask: gray colored areas indicate the time points at which classification did not survive the Bonferroni correction for multiple comparisons, while all other colors correspond to above-chance classification generalization.

## Further exploratory analyses of reliability and validity

### Split-half analyses

In a first set of exploratory split-half analyses, we examined whether, with half as much data, we could still reliably decode the content of children’s WM, and how the decoding generalized across time. Even when training/testing only within the early half, classification was robust during the majority of the Sensory period, and within the first 500ms of the Delay period, but not during the Baseline period (Figure 4A). Further, the same representations detected early in the Sensory period were observed throughout the Sensory and Delay periods (Figure 4A). The results for the late-half showed that we reliably classified data during the Sensory period and most of the Delay period (though not consecutively), but not during the Baseline period (Figure 4B). Here too, the representations detected early in the Sensory period were present throughout the Sensory and Delay periods (Figure 4B). We also investigated the consistency or stability of the representational structures in WM throughout the entire session, by conducting cross-decoding analyses between the early and late halves. We observed reliable cross-decoding during most of the Sensory period and the first 1000ms of the Delay period (though not consecutively). Finally, here too, representations detected early in the Sensory period were present throughout the Sensory and Delay periods (Figure 4C). This suggests that representational structures detected in the early half of the testing session were comparable to those detected in the late half of the testing session. In sum, these split-half classification results show that our approach detects robust representations, which are furthermore stable across the testing session, lending support to the overall reliability and validity of the approach.

**Figure 4.**
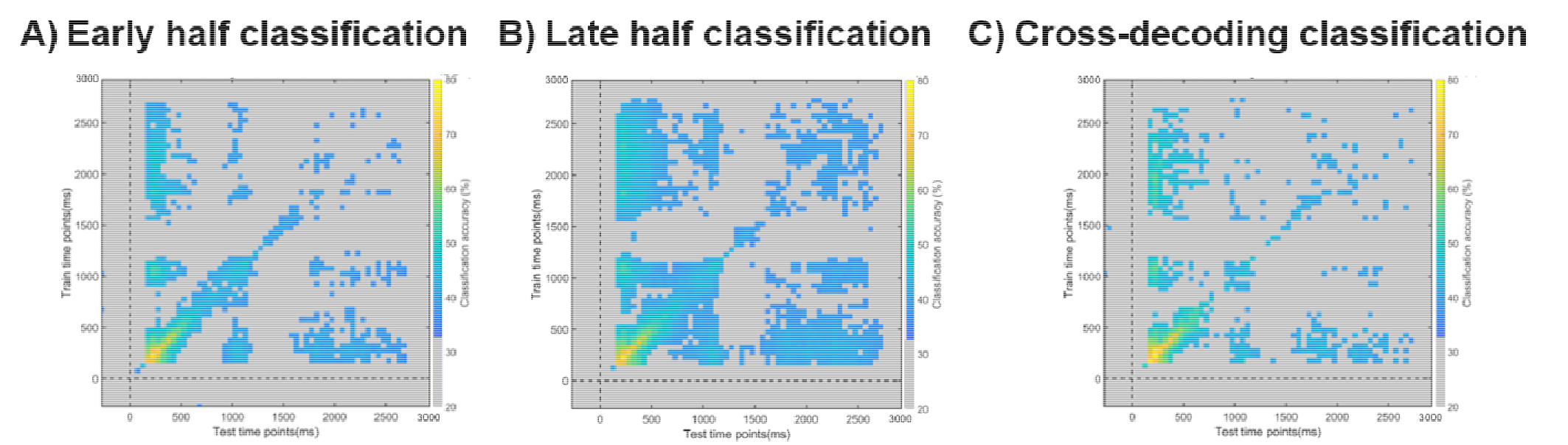
Classification performance on EEG data split into early (‘blocks’ 1-4) and late (‘blocks’ 5-8) halves. A) Classification results in the early half of the testing session, B) Classification results in the late half of the testing session and C) Cross-decoding results between the early and late halves (training the classifier on early half data, testing the classifier on late half data). Across all panels, Average classification values are overlaid by a significance mask: gray fields indicate the time points at which classification did not survive the Bonferroni correction for multiple comparisons, while all other colors correspond to above-chance classification generalization.

### Category confusability

We next examined the classification performance of specific categories, to identify which stimulus categories were most confusable, using the confusion matrices from the time-average classification (Figure 5). First, during the Baseline period, all categories at test were almost equally confusable with each other, since all of the values were near chance (33%). Next, during the Sensory period, we observed that verbal information was most accurately decoded (71%) and least confusable with either visual or spatial information. Whereas visual (52%) and spatial (51%) information were more confusable with each other (35%). The confusability patterns in the Sensory period were echoed during the Delay period, though with less pronounced distinctions between the categories, consistent with slightly lower overall decoding accuracy at Delay than at Sensory. These results show that each category was distinct enough from other categories, and that the specific pattern of differences between categories was consistent with theoretical distinctions between WM processes.

**Figure 5.**
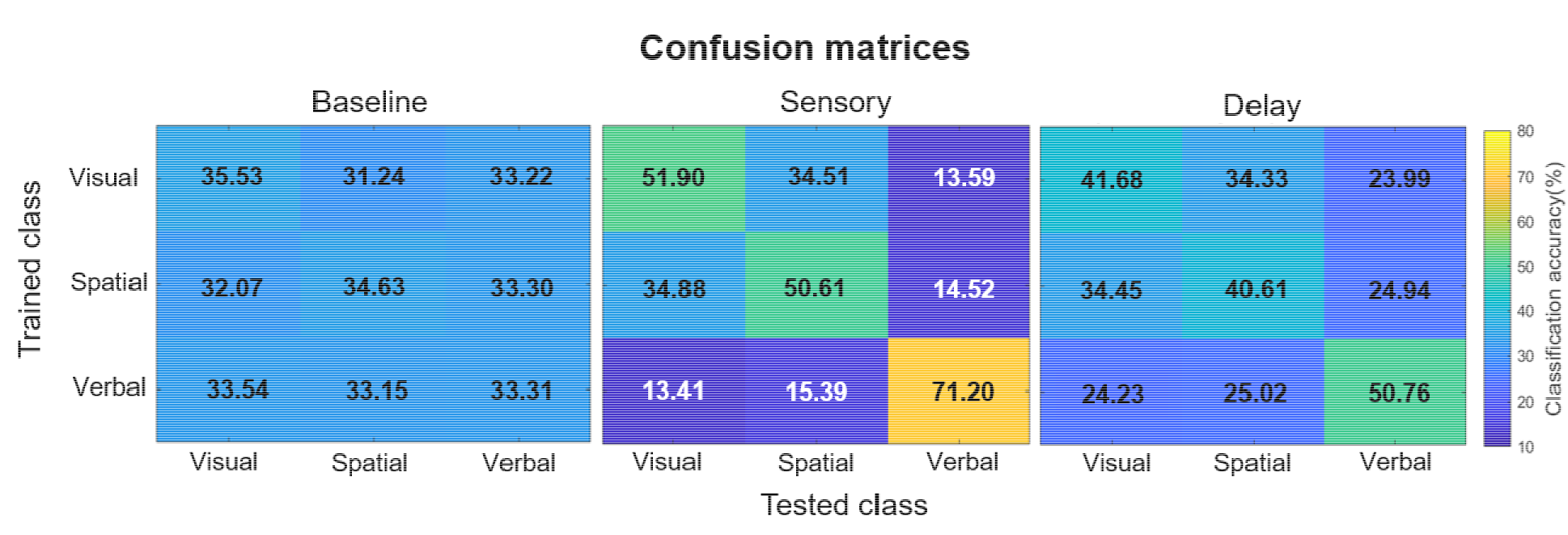
Confusion matrices for the three categories of information (visual, spatial, and verbal) during the Baseline, Sensory, and Delay periods. Training class labels are on the y-axis, and testing class labels on the x-axis. The tables show the proportion of trials where a category was confused with any of the other three categories.

### Individual differences in multivariate decoding

When it comes to individual classification performance, we had two main assumptions with regards to the consistency of our measures across participants. First, we assumed that classification performance would be higher during the Sensory period than during the Delay period. Second, we assumed that individuals with higher classification performance at Sensory will also have higher classification performance at. To test these assumptions, we first compared average classification accuracy at Sensory and Delay using a paired-sample right-tailed t-test. Second, we quantified the strength of the relationship between classification accuracy at Sensory and Delay via Pearson correlation.

The results of the paired-sample right-tailed t-test and Pearson correlation confirmed our assumptions (Figure 6). Namely, classification accuracy was higher during the Sensory period than in the Delay period across participants (*t*(19) = 9.87, *p* = 3.23e-09), by an average of 13.5%. Moreover, classification performance was highly correlated across individuals (*r*(18) = .84, *p* < 0.001). That is, most participants with high decoding accuracy in the Sensory period also tended to have high decoding accuracy during the Delay period. This suggests that our results were driven by patterns present across the whole sample, and that our approach is consistent across individuals since decoding accuracy across individuals was consistent across the Sensory and Delay periods.

**Figure 6.**
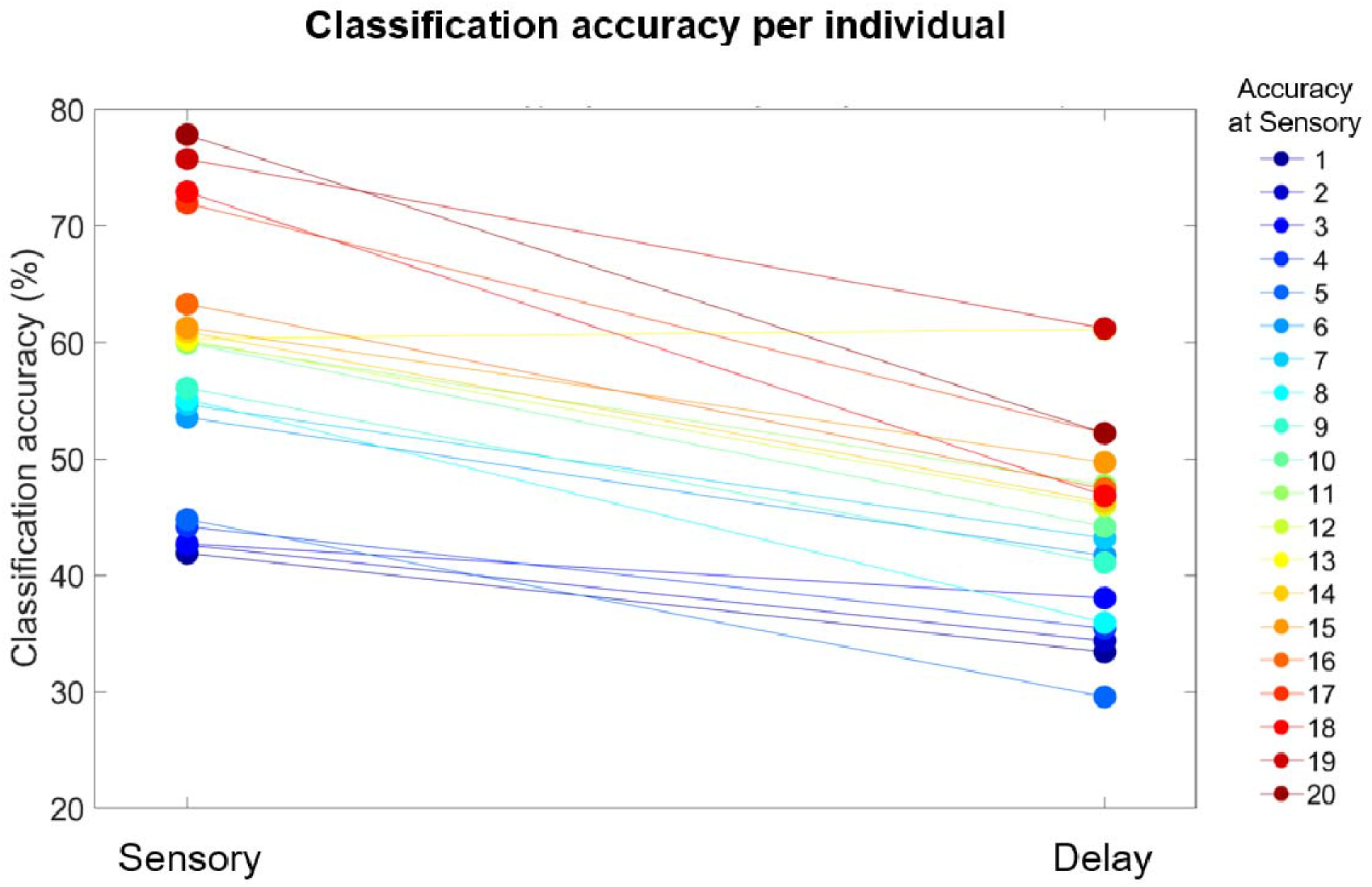
Individual differences in classification accuracy. Each dot depicts the average decoding accuracy from a participant during the Sensory and Delay periods, and dots from the same individual are connected by lines. The colors are organized according to the decoding accuracy during the Sensory period, from lowest accuracy (blue) to highest accuracy (red).

### Decoding across age

Figure 7 shows decoding accuracy as a function of age. One can immediately see that successful decoding during Sensory and Delay periods was present at all ages, from the youngest to the oldest participant. Moreover, regression analyses on decoding accuracy as a function of age did not reveal any significant relationship between age and decoding accuracy in any period of the trials: Baseline (*R^2^* = 0.03, *p* = 0.46), Sensory (*R^2^* = 0.02, *p* = 0.57), and Delay (*R^2^* = 0.03, *p* = 0.51). As Figure 7 indeed shows, in each of the periods of interest, decoding accuracy was stable across the entire age range. Our above-chance decoding thus does not seem to be driven by high-performing older children.

**Figure 7.**
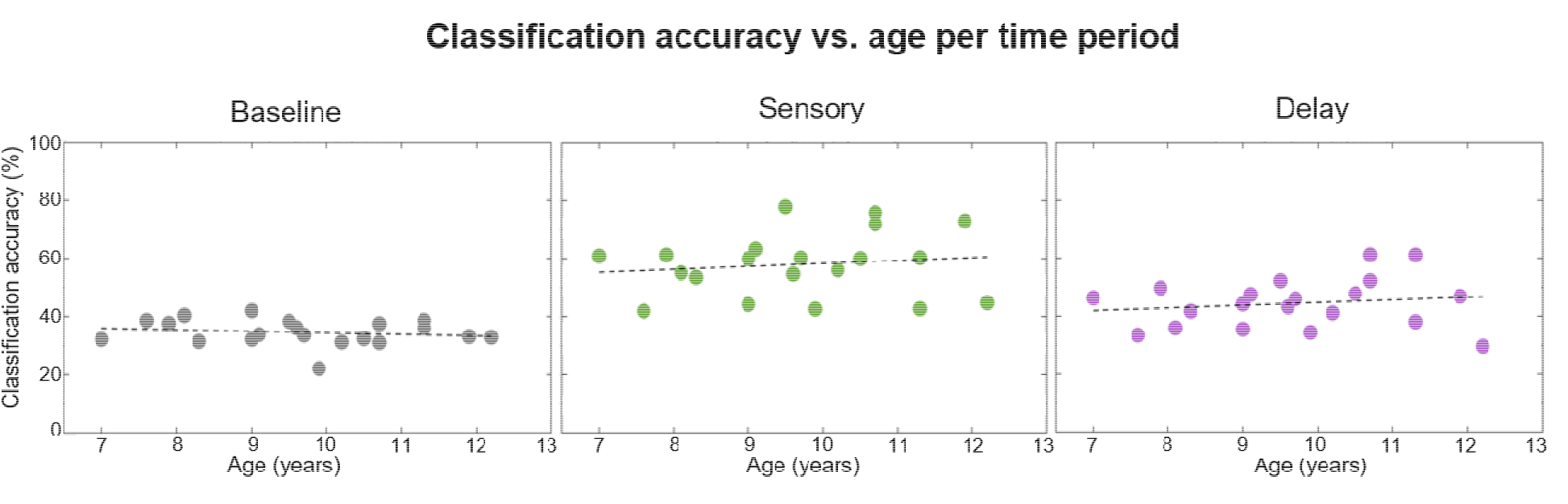
Decoding accuracy as a function of age. For each period (Baseline, Sensory, and Delay) we examined whether there was a relationship between classification accuracy and age. The age of the participant is depicted along the x-axis, and the classification accuracy is along the y-axis. Each participant is represented by one dot in each plot. The black dashed regression line on each panel shows the relationship between age and classification accuracy, which was not reliable for any of the three periods.

As to whether our decoding results may have been driven by younger children struggling to process the verbal stimuli, a BANCOVA showed that the best model, i.e., the model that best fit the data included the main effects of Category and Age. However, removing the covariate of Age from the best model only made the best model 2.5 times worse. This is a level of evidence that falls within the anecdotal range (Jeffreys, 1961; Kass & Raftery, 1995; Schönbrodt et al., 2017), suggesting that Age did not have an important effect in the best model, and that behavioral accuracy was not meaningfully affected by age. This result is not unexpected given that the participants only had to maintain one item, which children in the age range of our participants can easily do (Riggs et al., 2006). Next, the youngest participants’ behavioral accuracies per category were 81.25% for visual, 79,53% for spatial, and 83,59% for verbal, compared to the overall sample’s 90.78% for visual, 88.91% for spatial, and 90.59% for verbal As expected, the youngest participants had lower overall accuracy compared to the entire sample. However, their accuracy was lowest for the spatial category, not the verbal category, mirroring the entire sample. Further, their accuracy reduction compared to the overall sample was the smallest for the verbal category. Thus, in terms of behavioral WM performance, there was no evidence that the youngest children processed the nonwords in a drastically different way compared to the overall sample.

Finally, a comparison of confusion matrices between the entire sample and the youngest participants (Figure 8) revealed lower classification accuracies for the verbal category at Sensory and Delay for the youngest participants. That said, their classification accuracies were still high, and certainly above the 33% chance level: 61.69% at Sensory and 44.03% at Delay. These results further support that the youngest participants could successfully process nonword stimuli. It is thus highly likely that our results were driven by patterns present across the whole sample, regardless of age, lending evidence to the consistency of our approach across individuals.

**Figure 8.**
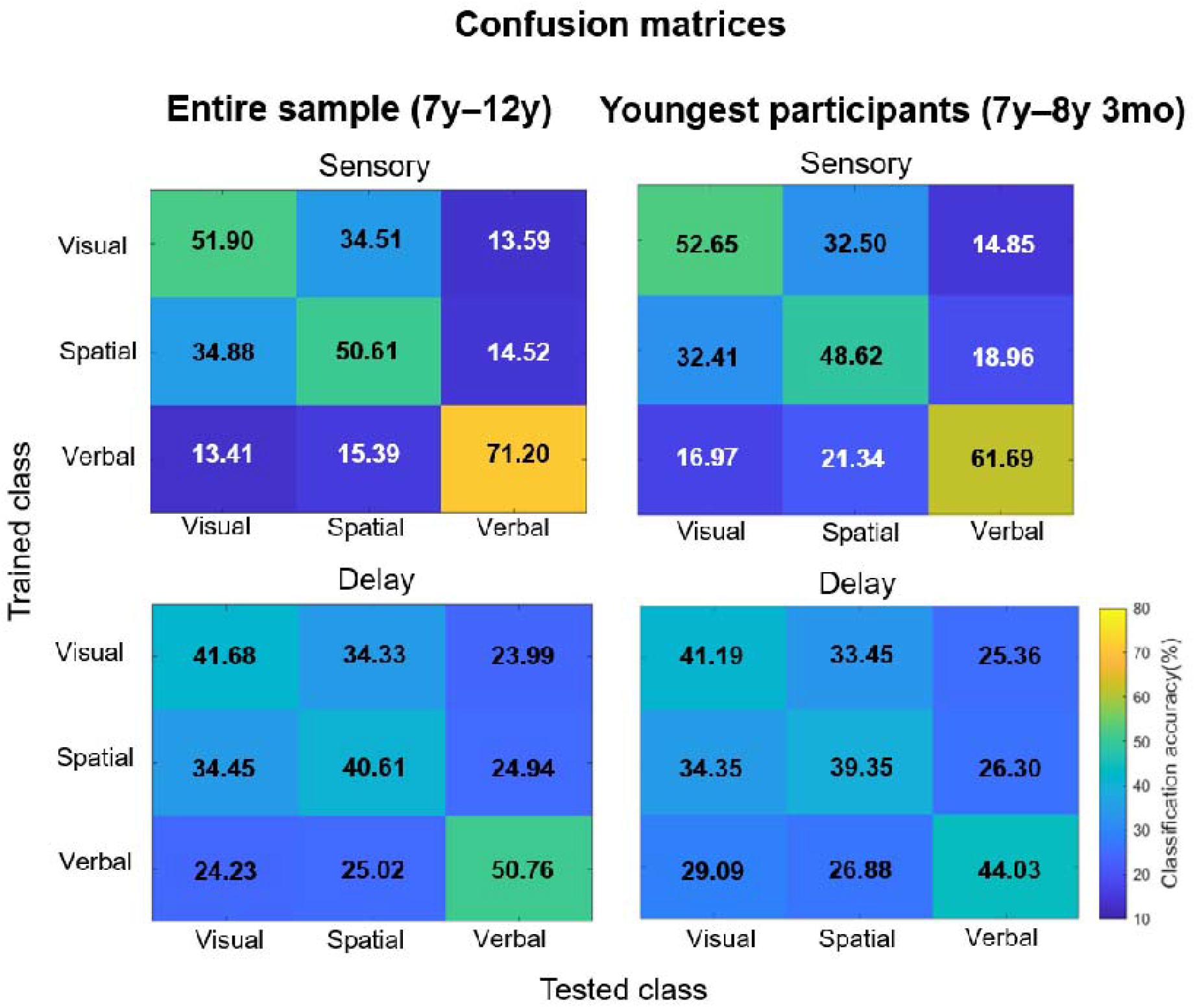
Confusion matrices showing classification accuracy for the three information categories (visual, spatial, and verbal) during the Sensory, and Delay periods. The results of the entire sample (repeated from Figure 5) are shown on the left, and the results of the 5 youngest participants (aged 7 years to 8 years and 3 months) are shown on the right, for comparison purposes. Training class labels are on the y-axis, and testing class labels on the x-axis. All results are shown on the same scale as in the bottom right corner. The tables show the proportion of trials where a category was confused with any of the other three categories.

## Discussion

In the present study, we aimed to deploy an analysis approach based on MVPA of EEG data to directly measure uninstructed WM representational content in children. Therefore, we developed a game-like computerized task with no maintenance instructions. We collected EEG data from children while they performed this task, and analyzed this data with multivariate pattern analysis methods. We reliably decoded the category of information during a WM maintenance period. Furthermore, we examined the temporal generalization of classification performance during the maintenance period and the consistency of the representations from sensory to delay periods. In addition, we conducted exploratory analyses of the reliability and validity of our results.

### Decoding the content of children’s WM

Although this is the first study to use MVPA on EEG signals to investigate children’s WM maintained memoranda, the results are comparable with the existing literature in adults. Namely, our delay period decoding accuracy between visual, spatial, and verbal categories in children’s EEG (44%) was comparable to decoding accuracy between visual, phonological, and semantic categories in young adults’ EEG (45.3%) in the Phase 1 task of LaRocque and colleagues (2013). This is especially impressive given that our sample consisted of children, a population whose WM performance is known to be lower than that of young adults (e.g., Gathercole, 1999; Gathercole et al., 2004; Salthouse, 1994), and whose EEG signals are not perfectly comparable to those of adults, due to physiological differences, higher spectral power in children than adults, etc. (e.g., Barriga-Paulino et al., 2011; Scerif et al., 2006). It is equally noteworthy that such results were obtained from an MVPA pipeline based on adult applications (e.g., the mini-block approach by Adam et al., 2020). Thus, even before delving into reliability and validity, these results already suggest that our approach is appropriate for decoding children’s WM content.

Inspired by methodological issues in measuring children’s maintenance mechanisms behaviorally, and the resultant theoretical confusion surrounding children’s ability to apply certain maintenance mechanisms spontaneously (e.g., in the case of refreshing and organization), we aimed to use an analysis approach, known to decode memory representations in adults, to tap into the content of children’s WM. We explored whether it is possible to infer children’s WM contents during maintenance, as a first step towards potential future studies on the maintenance mechanisms and strategies they use. As such, the results of this proof-of-concept study cannot in and of themselves reveal anything about strategies or maintenance processes and thus, for example, cannot answer whether, or at which age, children can spontaneously refresh information. However, since the approach appears sensitive and appropriate for detecting children’s WM representations, just like adults’ WM representations (e.g., Christophel et al., 2018; Lewis-Peacock et al., 2012, 2015; Rose et al., 2016), it could potentially be used as a base for addressing which maintenance processes children spontaneously use and when, as well as other open questions in the field of WM development.

### Exploring the reliability and validity of our approach

Given the topic of this special issue, although our paradigm was not optimized for such analyses, and although such analyses are not typically conducted (or at least not always reported) in the adult literature, we carried out exploratory analyses of reliability and validity to assess the use of our approach as part of a derivation chain in the field of WM. Within such constraints, these exploratory analyses nonetheless added further support to the main results.

Splitting the data into an early and late half showed two main lines of support for the consistency of the decoding results over time. First, the pattern of decoding accuracy in both the early and late halves was comparable to that of the overall pattern in the main analyses. Second, training the classifier on the early half data and testing it on the late half data showed that representational structures detected in the early half were similar to those in the late half. Both of these results suggest that, despite potential fatigue over the course of an experimental session, decoding is nonetheless reliable, and that WM representations remain similar over the course of a testing session.

Given our task design, we expected decoding accuracy across the different stages of WM to follow a given pattern. At Baseline, we expected decoding not to rise above chance levels, as this period presumably mainly contained noise. Likewise, there was no blocking of trials by category, such that anticipatory category-related activity would be detected during this period. Next, the Sensory period involved the perception, recognition, and encoding of the information to be remembered. Since this information was physically present on the screen during this time, we expected decoding accuracy to be the highest during the Sensory period. Finally, since the Delay period involved the maintenance of information in WM in the absence of the to-be-remembered stimuli on the screen, we still expected above-chance decoding accuracy, though lower than in the Sensory period. These patterns were borne out by time-average results, showing that our decoding procedure correctly responded to 1) noise at Baseline, 2) differences between observed categories of stimuli at Sensory, and 3) differences between representations of maintained categories of stimuli at Delay.

Based on classic WM theoretical accounts whereby WM is split into separate visuo-spatial and verbal domains (Baddeley & Hitch, 1974; Baddeley & Logie, 1999), if our approach truly captured WM content, the results would reveal verbal information to be distinct from visual and spatial information, while the latter two would be more confusable. To assess this, we examined patterns in the confusion matrices generated as part of the time-average analyses. We specifically observed that information labelled as verbal at training was the least confusable with information labelled as visual or as spatial at test, and that information labelled as visual and spatial at training was relatively confusable with each other at test. This is in line with classic theories of domain-specificity for visuo-spatial and verbal information (Baddeley & Hitch, 1974; Baddeley & Logie, 1999), and suggests that our approach measured actual maintained WM content.

The last two analyses confirmed that our results were not driven by a subset of participants, but by consistent values across the entire sample. Namely, we observed that the majority of participants had slightly higher decoding at Sensory than at Delay, and that those individuals that had high decoding accuracy at Sensory also had high decoding accuracy at Delay. Several explanations exist as to why some individuals had higher decoding at both stages than others. Apart from differences in WM ability (here, how well one can represent physically absent information), differences in electrode preparation (e.g., the amount of electrolyte used) and skull thickness could also have driven individual differences in decoding. Though we collected no data at this stage to explore any such possible accounts, this may be an interesting direction for future research. Skull thickness is known to change with age (e.g., Calderbank et al., 2016), so this variable could have potentially driven any changes in decoding accuracy with age. However, there was no evidence for decoding changes as a function of age, as classification accuracies were evenly distributed across participants regardless of age. Importantly, there was no evidence for the observed decoding results being driven either by high-performing older participants, or by younger participants’ potential differences in processing the verbal stimuli. According to classic theories of reading development (e.g., Chall, 1983), younger children have different strategies for processing verbal stimuli than do older children. Thus, it could be possible that younger children had different (slower/less adaptive) ways to process and maintain our verbal stimuli, which could have affected our decoding results. We did observe lower classification accuracy, i.e., greater confusability, for the verbal category in the Sensory and Delay periods in younger participants compared to the overall sample. However, their classification accuracy was still well above chance, and there was no associated behavioral performance decrease. Thus, rather than not having adequate strategies for processing and maintaining the nonword stimuli, the younger children could have relied a bit more on the visual characteristics of the nonwords to maintain them in WM. Either way, their behavioral and decoding results did not indicate any major differences from the overall sample. Taken together, these results suggest that our approach is reliable across time within a testing session, across individuals. Further, they suggest that our approach measured what we set out to measure, that is, differences in the content of children’s WM.

### Limitations

An inherent limitation of our study was that its design and resource allocation were optimized for the main analyses, and aim to provide a proof of concept of decoding children’s WM content, but not for our analyses of reliability and validity. For instance, our task was designed to be easy to complete successfully across the 7-12 age range. In addition, our sample size (n=20) was appropriate for a proof-of-concept study, but relatively small for regression analyses (20 participants). Now that our proof of concept appears to have been successful, future studies should include retests and replications in larger samples.

Another potential issue with our design lies in its lack of masks, i.e., irrelevant stimuli that would overwrite the contents of sensory memory, and prevent such sensory representations from driving the decoding results. We intentionally refrained from adding any additional information other than the to-be-remembered items and their probes, both to keep the task as simple as possible, and to prevent any potential biasing effects resulting from masks (see “attractive bias” in Lorenc et al., 2021). A point that is rarely acknowledged in the use of masks is that they can function as external distractors (i.e., irrelevant stimuli that should not be attended; reviewed in Rademaker et al., 2015, p. 1-2). Since children are known to be more susceptible to visual distraction than adults (e.g., Plude et al., 1994; Trick & Enns, 1998), we did not wish to risk replacing sensory memory effects with potential distractor effects. This point notwithstanding, our current design does not let us directly assess whether representational content at maintenance consists merely of a persisting sensory trace. Indeed, there are some differences in features across categories that could have driven the decoding. For example, the verbal stimuli occupied less vertical space on the screen than the visual and spatial stimuli, or that the visual stimuli were more colorful than the other stimuli (Supplemental Figure 1). However, that would be unlikely, since the perceptual and informational content of a visual stimulus only seem to persist for a maximum of 500ms after stimulus offset (Irwin & Yeomans, 1986; Massaro & Loftus, 1996; Sperling, 1960), and we found significant decoding for almost the entire Delay period, consistent with comparisons of sensory memory and WM such as Cappiello & Zhang, 2016). Similar differences between verbal and visual stimuli did not deter LaRocque et al. (2013), whose paradigm we leaned on to construct ours, from making conclusions about WM maintenance. Finally, in an exploratory searchlight analysis (detailed in the Supplementary Materials; see Supplemental Figure 5), we found above-chance decoding accuracy across the entire electrode montage, which would be unlikely in the event that only purely sensory-feature differences in memoranda drove the classification. Though our current paradigm cannot undeniably rule out potential influences of sensory features in the decoding, observing above-chance classification in regions with limited perceptual representations, and during periods that extend far beyond stimulus presentation shows that our decoding analysis did capture maintained representational content in WM.

Finally, we used a relatively strong highpass filter of 1Hz coupled with baseline correction to clean our EEG data before classification. Highpass filters as low as 0.05Hz have recently been recognized to produce decoding artefacts, particularly in temporal generalization, by temporally shifting the signal and generating spurious activity in the baseline period, which baseline correction then further exacerbates by shifting it onto the signal later on in the trial (van Driel et al., 2021). Our highpass filter choice was motivated by the need to perform accurate ICA on our EEG data (e.g., Winkler et al., 2015) which was necessary as children’s data are typically noisier than that of adult data. To test whether our above-chance decoding could have been a result of such issues, we conducted a check recommended in van Driel et al. (2021, p.16; see Supplementary materials: Supplemental preprocessing checks). This indicated that it is highly unlikely that our decoding results were contaminated by artefacts resulting from our preprocessing pipeline. Nonetheless, future studies employing MVPA on EEG data may wish to verify their filter settings before conducting classification analyses to avoid potential pitfalls as highlighted by van Driel and colleagues (2021).

## Conclusion

The current study demonstrates that children’s WM contents can be decoded using EEG MVPA techniques established in adults together with a simple behavioral paradigm, in a manner that is promising in terms of reliability and validity. Though the main insights the study provides are what children are maintaining rather than how, the framework we developed as part of this study can serve as a base for investigations of maintenance mechanisms, or questions related to representational content. As such, this study provides a much-needed stepping stone for strengthening the derivation chain in the field of WM development.

## Acknowledgments

We would like to thank all the families that participated in the experimental sessions, and Tom Hilbert for his inputs on assessing the functionality of our filter. This work was completed with support from the Swiss National Science Foundation to Evie Vergauwe [Grant PCEFP1_181141], from the Jacobs Foundation to Nora Turoman [Grant 2021-1417-00], and from the NIH to Megan deBettencourt [5F32MH115597 and K99 MH128893].

