## Supplementary materials for "Decoding the content of working memory in school-aged children"

**Supplemental Figure 1: All Stimuli together**

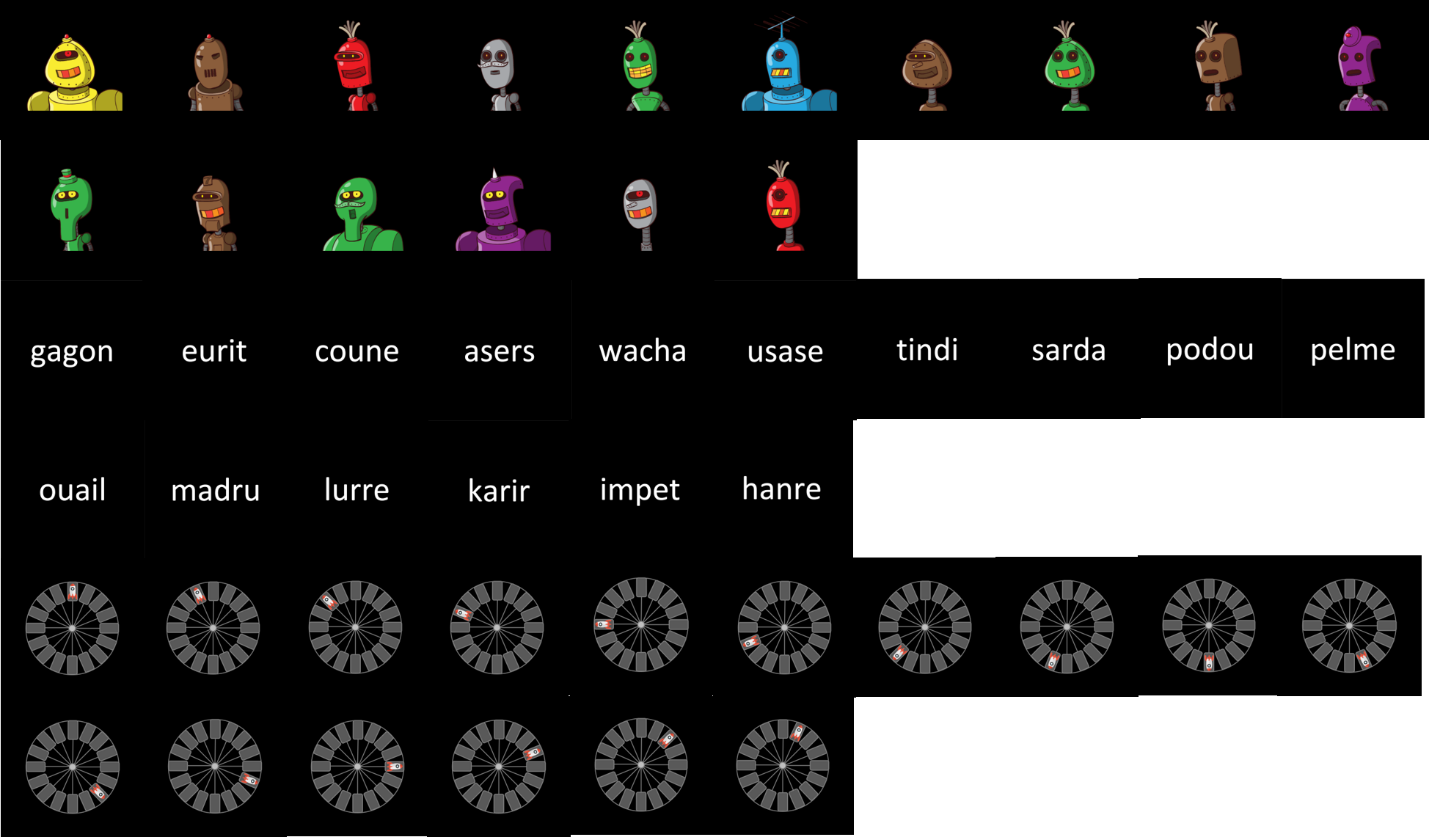

**Supplemental Figure 1.** All stimuli from all categories (visual on top, verbal in the middle, and spatial on the bottom) displayed together, for comparison purposes. All stimuli subtended 5 degrees of visual angle during their presentation.

**Supplemental EEG classification results: Temporal generalization results using 20ms time-bin averages**

A classifier was trained and tested on data from 20ms time bins steps over a trial (without the Baseline period, i.e. 0 to 3000ms), where EEG voltage data were averaged over each time bin. A temporal generalization classification examined the unfolding of information category representations across time. Consistent with the 50ms time bin results reported in the main text, we found that representations that were detected during the early Sensory period could be detected over the rest of the Sensory period, and the Delay period. This is reflected in three patterns per Supplementary Figure 2: 1) the high accuracy levels on the main diagonal on the graph, 2) data trained on Delay period representations showing higher accuracy when tested on early Sensory period data, and 3) data trained on early Sensory period representations showing somewhat high accuracy when tested on Delay period data. The results using 20ms time bins thus corroborate the results of the 50ms time bin analysis.

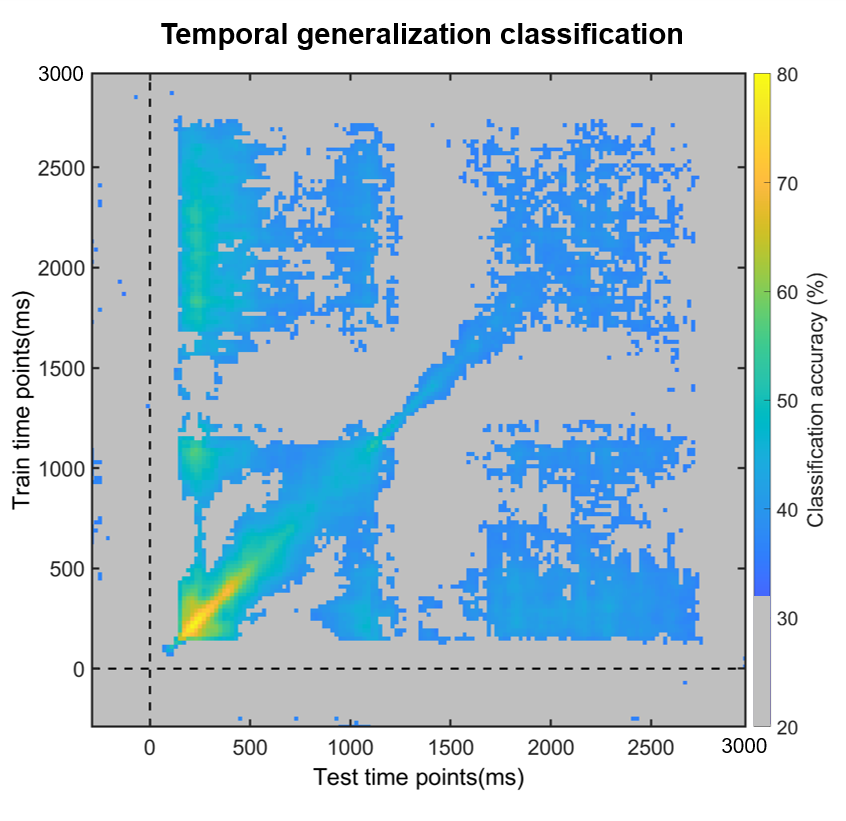

**Supplemental Figure 2.** Results of the temporal generalization analysis, where a classifier was trained on one time point and tested on all other time points. Here, 20ms time-windows were used in the classification. Average classification values are overlaid by a significance mask: gray fields indicate the time points at which classification did not survive the Bonferroni correction for multiple comparisons. All other colors correspond to classification generalization per the bar on the left of the figure.

**Supplemental exploratory reliability and validity results: Split-half analyses using 20ms time-bin averages**

As in the 50ms time bin analyses reported in the main text, datasets were split into early (‘blocks’ 1 – 4 in the testing session) and late (‘blocks’ 5 – 8) halves, on which the same classification schemes as before were applied separately. Even when training/testing only within the early half (Supplementary Figure 3A), or only within the late half (Supplementary Figure 3B), classification was robust during almost the whole Sensory and Delay period, with no above-chance decoding during the Baseline period. Representations detected early in the Sensory period were present throughout the Sensory and Delay periods. Thus, our method can reliably detect representations in a stable manner across the testing session, even when provided access to only half of the data.

To investigate the stability of representational structures from the beginning to the end of the session, we conducted a cross-decoding analysis of the early versus the late halves. With 20ms time bins, just as for 50ms time bins, we observed reliable cross-decoding during the entire Sensory and first 1000ms of the Delay period (Supplementary Figure 3C). This suggests that representational structures detected in the early half of the testing session generalize for a portion of the late half of the testing session.

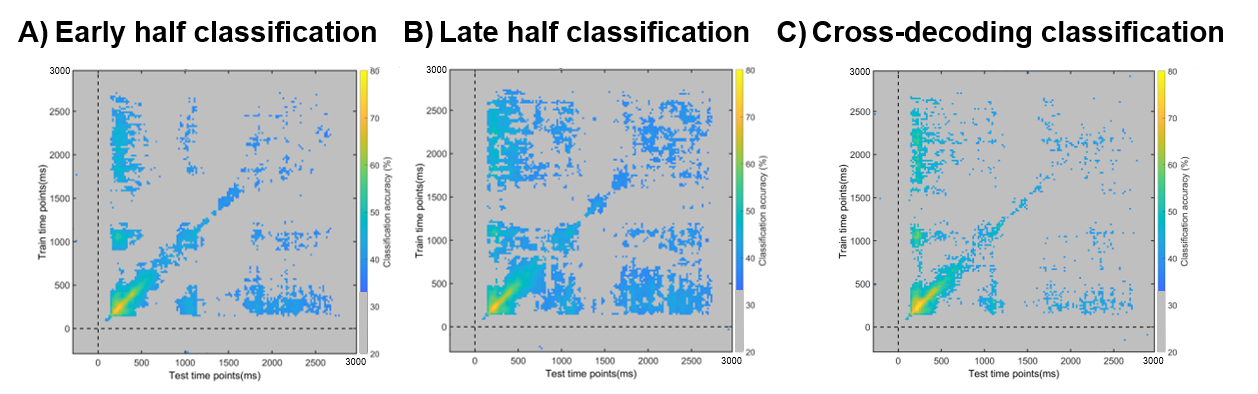

**Supplemental Figure 3.** Classification performance on EEG data separated into early (blocks 1-4) and late (blocks 5-8) halves. A) shows early half classification results, B) shows late half classification results, and C) shows cross-decoding results between the early and late halves. Across all panels, average classification values are overlaid by a significance mask: gray fields indicate the time points at which classification did not survive the Bonferroni correction for multiple comparisons. All other colors correspond to classification generalization per the bar on the left of the figure.

**Supplemental Figure 4:**

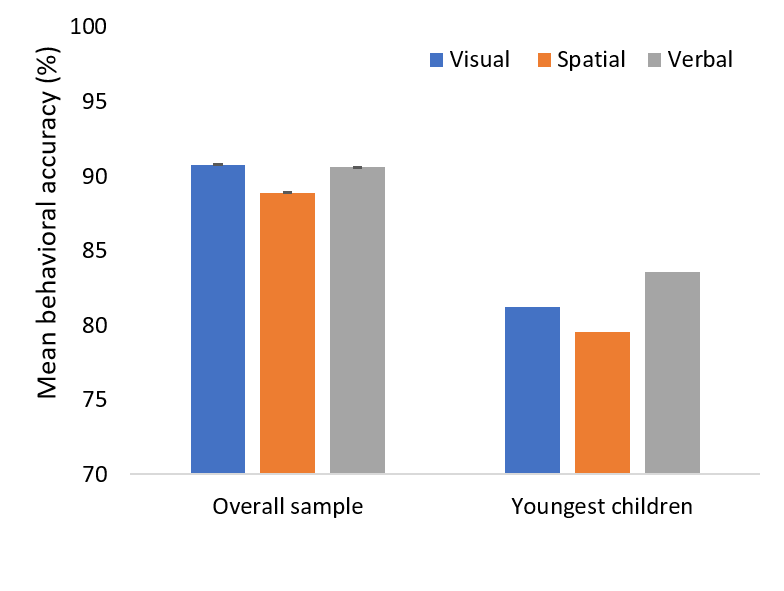

**Supplemental Figure 4.** Results of the comparison of behavioral accuracy per stimulus category (visual plotted in blue, spatial plotted in orange, and verbal plotted in gray) between the youngest children in the sample (5 children aged between 7 years and 8 years and 3 months), and the overall sample used in the study. Error bars represent the standard error of the mean, and are present on all bars, but are too small to be visible for the youngest children. The plot shows that, although accuracy is smaller overall for the youngest children than for the whole sample, the pattern across stimulus types is highly similar, with spatial stimuli having the lowest accuracy.

**Exploratory searchlight analysis**

To attempt to disentangle whether the decoding results over the Delay period were driven by differences in stimulus features across categories or genuine maintenance processes, we conducted an exploratory searchlight analysis, as suggested by a reviewer. To this end, we applied the same analysis approach as for the time-average analysis in the main manuscript, except that the features in the classifier were voltage amplitudes for each electrode and its nearest neighbors per 50ms time-bin. The results across time-bins were then combined and averaged to obtain a snapshot of classification accuracy across the Delay period.

The results (Supplementary Figure 5) show that although highest for posterior electrodes, classification accuracy was above theoretical chance (33%) across the electrode montage. Therefore, it is highly unlikely that stimulus feature differences alone drove the classification, as this would have presumably resulted in above-chance decoding only in posterior areas. That said, *maintenance* of sensory features could also have given rise to above-chance decoding in posterior areas (Awh & Jonides, 2001; Emrich et al., 2013; Serences et al., 2009), and the current paradigm has no way of disentangling the two. The current results could thus be reflecting memoranda being maintained in a distributed fashion across regions and levels of the entire cortex (Christophel et al., 2017; D’Esposito & Postle, 2015; Zimmer, 2008), and/or a combination of purely sensory-level influences or maintenance of sensory information together with more abstract maintenance strategies or even general cognitive control processes (e.g., D’Esposito & Postle, 2015; Emrich et al., 2013). Either way, observing above-chance decoding in regions with limited perceptual representations, and during time periods extending after stimulus presentation makes it highly unlikely that only sensory-feature differences across categories drove the decoding.

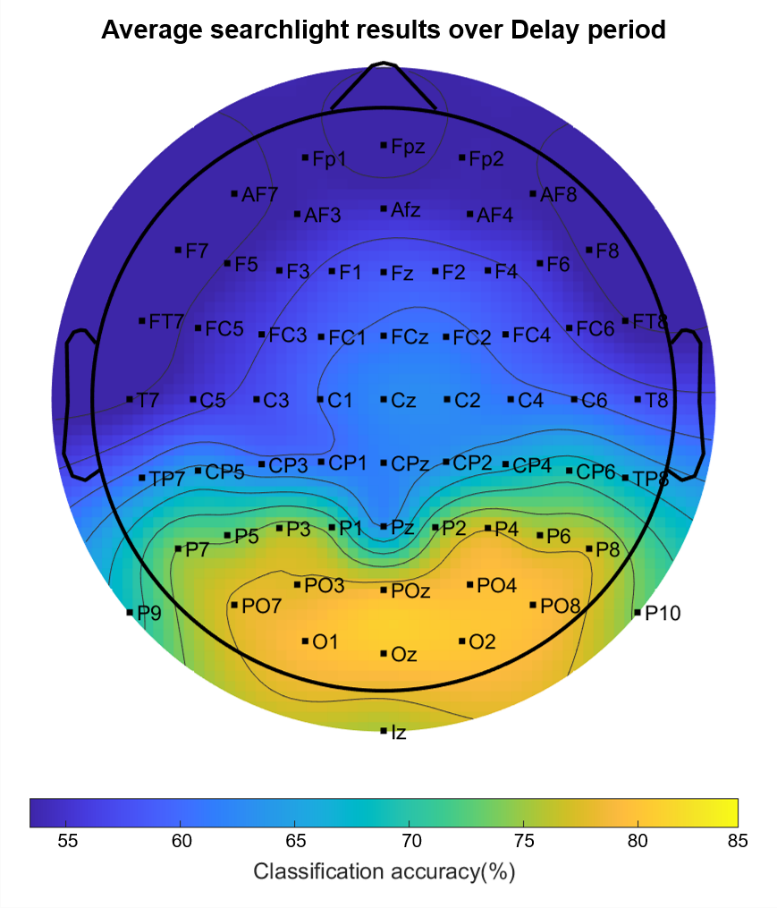

**Supplementary Figure 5.** Results of searchlight analysis showing average decoding accuracy across the entire Delay period (1000 – 3000ms). Decoding accuracy across the entire electrode montage was above theoretical chance (33%), which would have been represented in grey.

**Supplemental preprocessing checks**

To check if the bandpass filter (1-40Hz) that we used to clean the data before ICA and classification created any artefacts that could have skewed our decoding results per van Driel et al. (2021), we first verified whether the temporal generalization results show above-chance decoding before and after baseline correction (as recommended by van Driel et al., 2021, p.16). Below we show the results of this check, starting with Supplemental Figure 6.

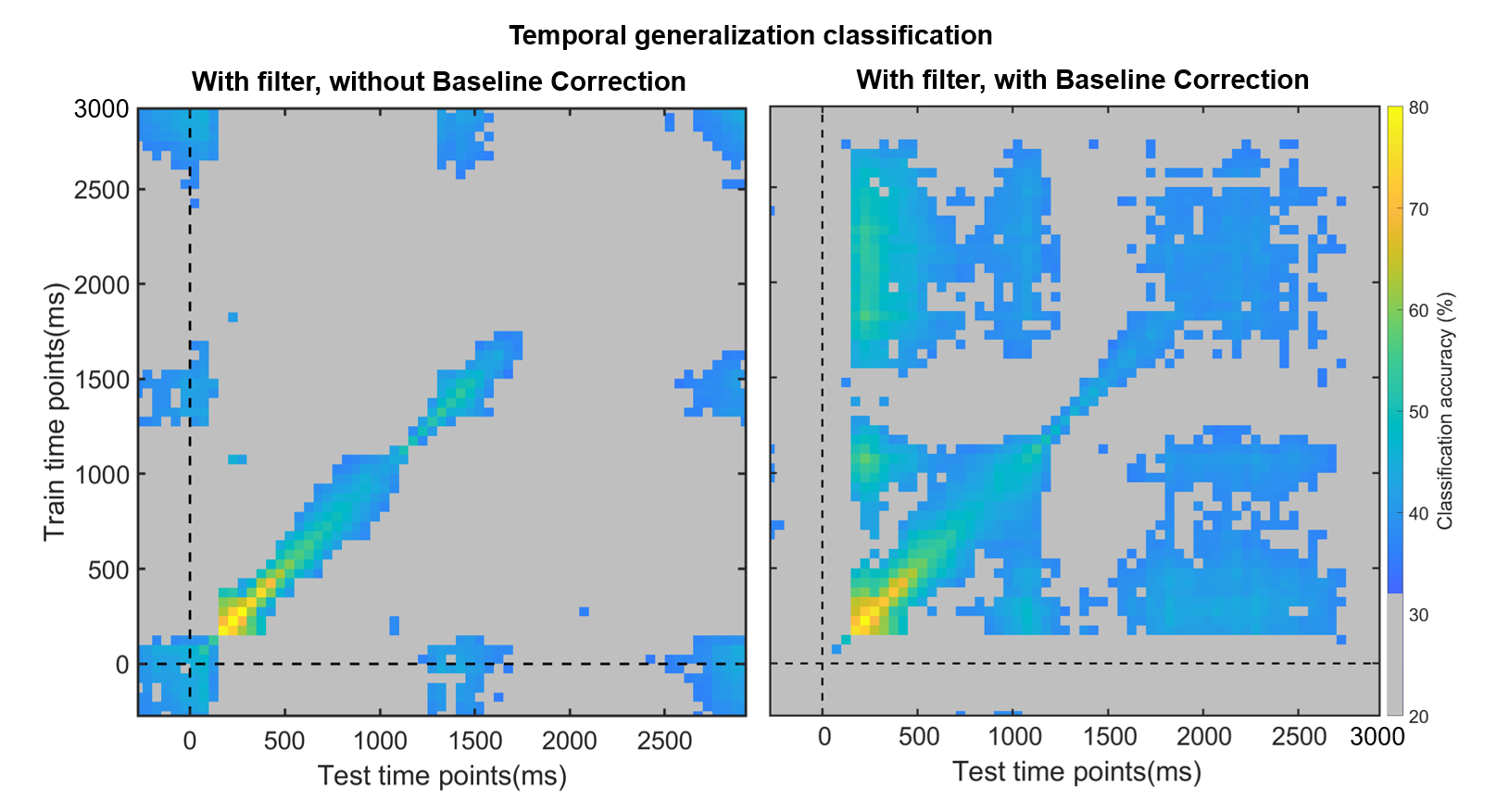

**Supplemental Figure 6.** Results of the first part of checking our EEG preprocessing pipeline for spurious filter-driven activity during the baseline period that may have become temporally shifted after baseline correction. The left side of the plot shows above-chance decoding during the baseline period after filtering, while the right side of the plot shows that this decoding has been eliminated after baseline correction. This stands in contrast with the van Driel et al. (2021) manuscript, which found baseline correction to further exacerbate above-chance decoding during the baseline period, as well as increase above-chance decoding during the rest of the trial. In these exploratory temporal generalization analyses, as above, a classifier was trained on one time point and tested on all other time points (50ms time-windows were used). Classification values are overlaid by a significance mask: gray fields indicate the time points at which classification did not survive the Bonferroni correction for multiple comparisons. All other colors correspond to classification generalization per the bar on the left of the figure.

Thus, as in the van Driel et al. (2021) manuscript, we saw above-chance generalization during the baseline period with filtering and without baseline correction. However, in contrast with the van Driel et al. (2021) manuscript, we did *not* find artefacts even with a 1Hz highpass filter and baseline correction. This led us to believe that, perhaps, our filter design did not actually temporally shift the signal to begin with. To explore this, we plotted the frequency and phase response of the Butterworth bandpass filter with cutoff frequencies at 1Hz and 40 Hz that we used (Supplemental Figure 7).

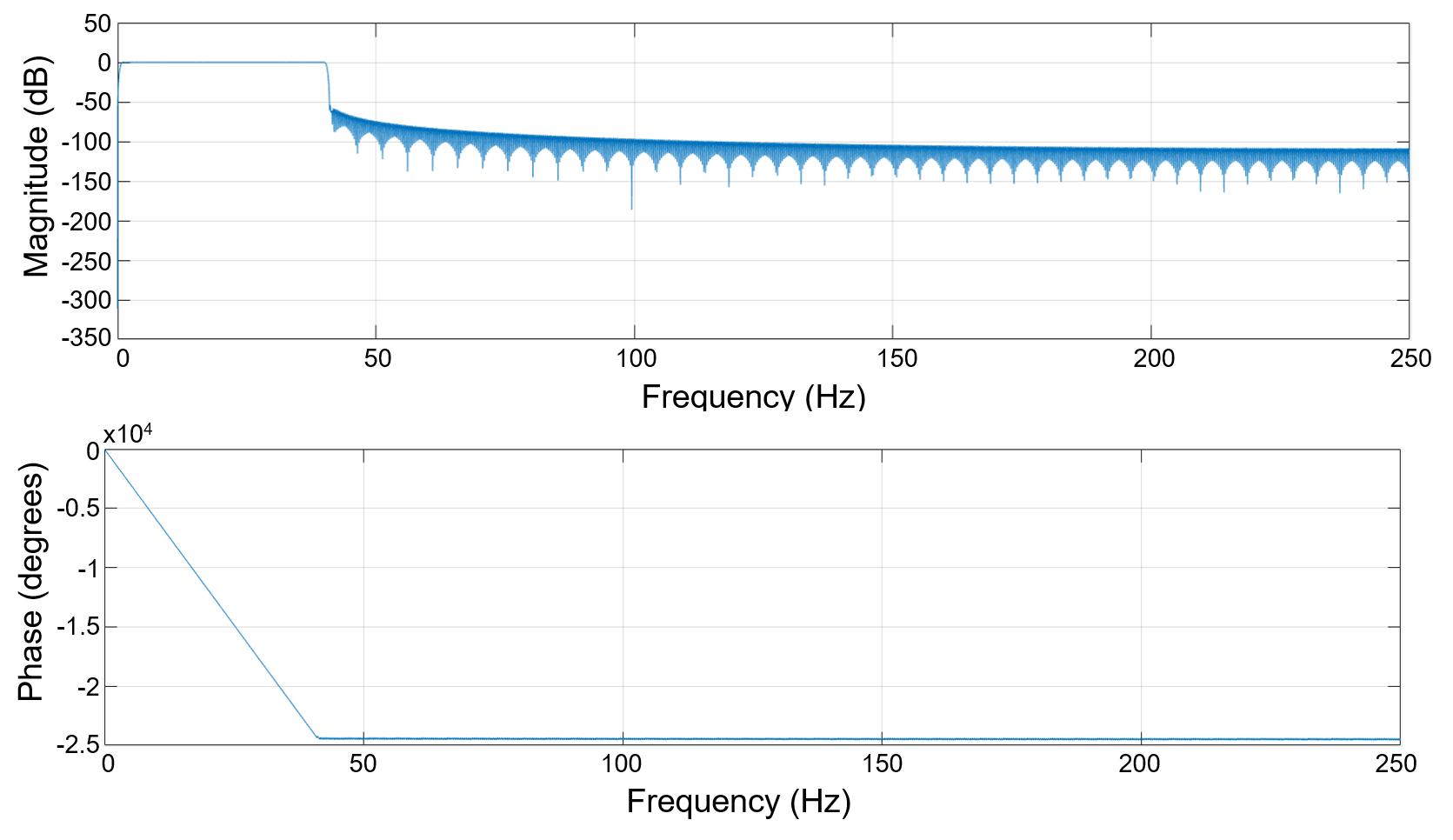

**Supplemental Figure 7.** Results of the second part of checking our EEG preprocessing pipeline. The top panel shows the frequency response, while the bottom panel shows the phase response of the bandpass filter that we used (1-40Hz).

In the phase response of the filter above, we observe a linear decrease in the phase within the passband. This means that, if there were a temporal shift, all frequencies within the passband would be equally affected. Next, we simulated EEG signals containing sine waves with different frequencies (0.5Hz, 10 Hz, 30 Hz, 50 Hz) to see whether we observe any temporal displacements using our 1-40Hz bandpass filter (Supplemental Figure 8).

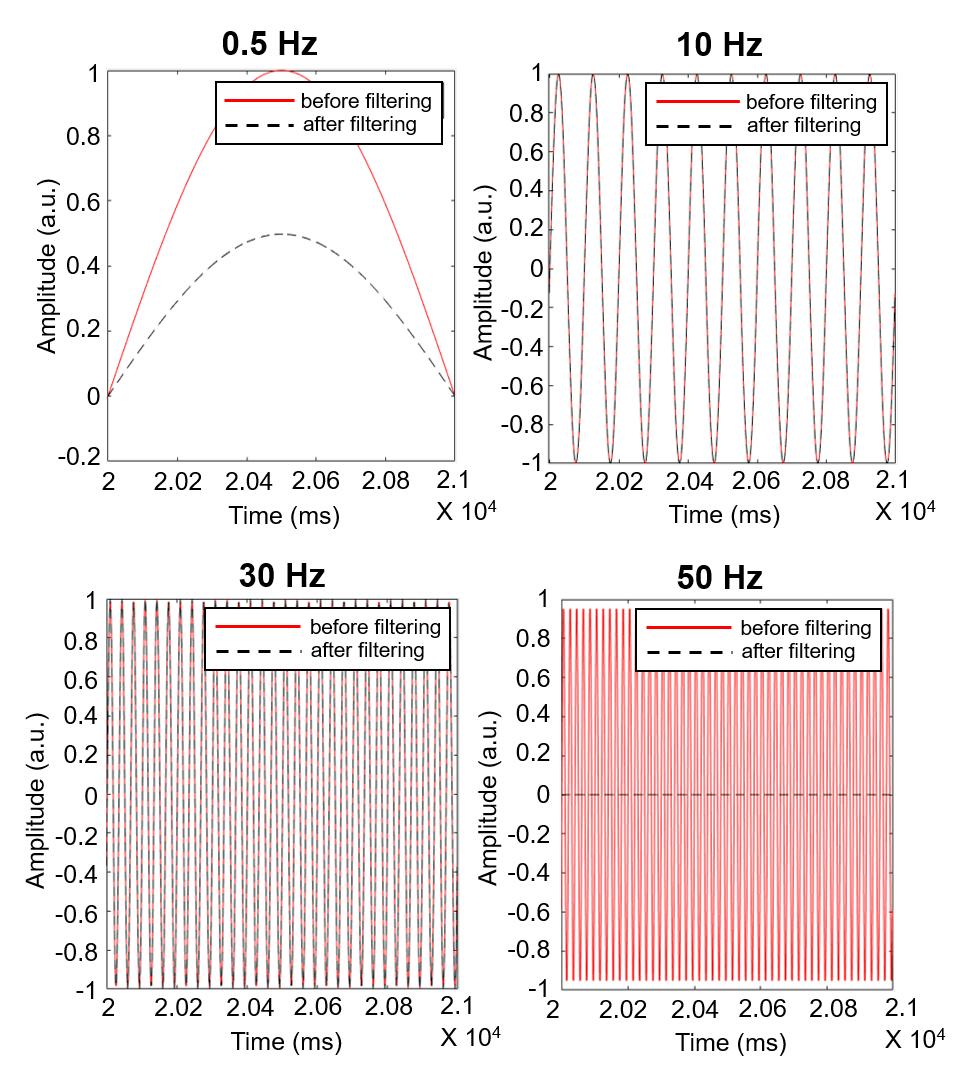

**Supplemental Figure 8.** Results of the third and final part of checking our EEG preprocessing pipeline. All four panels show the 1-40Hz filters being applied to simulated data at various frequencies: 0.5Hz, 10Hz, 30Hz, and 50Hz. The amplitude of the signal (in arbitrary units: a.u.) is plotted across an arbitrary time window from 2 to 2.1 x 10^4^ milliseconds. Data before filtering are shown in a black dashed line, while data after filtering are shown in a red continuous line. None of the data show shifts along the X axis, demonstrating that there were no temporal shifts as a result of the filter.

As expected, the filter fully removed the 50 Hz signal, and reduced the amplitude of the 0.5Hz signal without any temporal delays. For signals within the passband (10 Hz and 30 Hz), the filtered signal was identical to the original simulated sign wave, further showing that the filter that we used did not, and does not, introduce temporal shifts in EEG data.
